# *Candida auris*: multi-omics signature of an emerging and multidrug-resistant pathogen

**DOI:** 10.1101/528232

**Authors:** Daniel Zamith-Miranda, Heino M. Heyman, Levi G. Cleare, Sneha Couvillion, Geremy Clair, Erin Bredeweg, Attila Gacser, Leonardo Nimrichter, Ernesto S. Nakayasu, Joshua D. Nosanchuk

## Abstract

*Candida auris* is a recently described pathogenic fungus that is causing invasive outbreaks on all continents. The fungus is of high concern given the numbers of multidrug-resistant strains that have been isolated in distinct sites across the globe. The fact that its diagnosis is still problematic suggests that the spreading of the pathogen remains underestimated. Notably, the molecular mechanisms of virulence and antifungal resistance employed by this new species are largely unknown. In the present work, we compared two clinical isolates of *C. auris* with distinct drug susceptibility profiles and a *Candida albicans* reference strain using a multi-omics approach. Our results show that, despite the distinct drug-resistance profile, both *C. auris* strains appear to be very similar, albeit with a few notable differences. However, when compared to *C. albicans* both *C. auris* strains have major differences regarding their carbon utilization and downstream lipid and protein content, suggesting a multi-factorial mechanism of drug resistance. The molecular profile displayed by *C. auris* helps to explain the antifungal resistance and virulence phenotypes of this new emerging pathogen.

**Importance:** *Candida auris* was firstly described in Japan in 2009 and has now been the cause of significant outbreaks across the globe. The high number of isolates that are resistant to one or more antifungals, as well as the high mortality rates from patients with bloodstream infections, has caught the attention of the medical mycology, infectious disease and public health communities to this pathogenic fungus. In the current work, we performed a broad multi-omics approach on two clinical isolates isolated in New York, the most affected area in the USA and found that the omic profile of *C. auris* differs significantly from *C. albicans*. Besides our insights into *C. auris* carbon utilization and lipid and protein content, we believe that the availability of these data will enhance our ability to combat this rapidly emerging pathogenic yeast.

## Introduction

*Candida auris* is an emerging pathogenic fungus that was firstly described in 2009 after being isolated from the ear discharge of a patient in Tokyo, Japan (1). After the new species identification, a study in South Korea reported a misidentified *C. auris* strain isolated in 1996, which then became the first known case of human *C. auris* infection (2). Despite the fact that bloodstream infections are the main cause of mortality among *Candida spp* infections, *C. auris* strains have been isolated from various sites such as respiratory tract, bones, and central nervous system (3) as well as on a variety of abiotic surfaces (4), which suggests a metabolic plasticity to survive in distinct environments. The reports of *C. auris* outbreaks in all continents suggest that this pathogen is spreading rapidly across the globe and many of the isolated strains are resistant to at least one class of antifungals, or even multidrug-resistant (5-11). *C. auris* produces biofilms and can be very resilient in substrates commonly used in hospitals, features that are correlated with the frequency of reported hospital-associated infections as well as its increased resistance against antifungals (4, 9, 12-15). Additionally, its problematic identification suggests that reports regarding infection might be underestimated (16-18).

To understand the molecular mechanisms of infection, antifungal resistance and disease employed by this new pathogen, we performed a multi-omics approach using two clinical isolates of *C. auris* that were also compared to a standard *C. albicans* strain. The tested *C. auris* strains presented different levels of antifungal resistance, as one of them is highly resistant to fluconazole and slightly resistant to caspofungin. Both *C. auris* strains had very similar metabolic, lipid and protein profiles. However, both strains were significantly distinct when compared to *C. albicans*. Taken together our data show metabolic, lipidomic and proteomic similarities and differences between *C. auris* strains as well as in comparison with *C. albicans*, and our findings provide interesting insights into metabolic features, with some correlating with antifungal resistance.

## Methods

### Cell Lines

Two clinical isolates (MMC1 and MMC2) were acquired from Montefiore Medical Center (Bronx, NY, USA) under approved protocols in the Nosanchuk laboratory, and a standard *C. albicans* (ATCC #90028) strain was purchased from the ATCC. The strains were stored in -80 °C. Prior to use in experiments, cells were cultivated in YPD broth and seeded onto Sabouraud agar plates. For each experiment, one colony was inoculated in 10 mL of Sabouraud broth overnight at 30°C before use. Cells were transferred to 200 mL of fresh Sabouraud and incubated for additional 24 hours. After being extensively washed with PBS the cell pellets were frozen until the protein, metabolite and lipid extractions.

### Antifungal susceptibility

The antifungal susceptibility tests were carried out according to the CLSI protocol with minor modifications (19, 20). Yeast cells were inoculated in Sabouraud-agar for 48 hours at 30 °C and then stored at 4 °C up to one month for experimentation. One colony from each strain was inoculated in liquid Sabouraud and kept for 24 hours at 30°C under constant shaking. Cells were then washed in PBS and plated (2.5 x 103 cells/mL) in 96-well plates containing serial dilutions of amphotericin B, caspofungin and fluconazole. After 48 hours of incubation, cells were visually analyzed and the MIC was determined as the lowest concentration of a given drug that showed no apparent growth within all replicates.

### Proteomic analysis

Samples were submitted to metabolite, protein and lipid extraction (MPLEx) according to the protocol by Nakayasu *et al.* (21). Extracted proteins were digested with trypsin and resulting peptides were extracted with 1 mL Discovery C18 SPE columns (Supelco, Bellefonte, PA) as previously described (22). Digested peptides were suspended in water, quantified by BCA assay and 0.5 μg of peptides were loaded into trap column (4 cm x 100 μm ID packed in-house with 5 μm C18, Jupiter). Peptide separation was carried out an analytical column (70 cm x 75 µm ID packed with C18, 3 μm particles) using a gradient of acetonitrile/0.1% formic acid (solvent B) in water/0.1% formic acid (solvent A). The flow was set to 300 nL/min with 1% solvent B and kept for 15 min. Then concentration of solvent B was increased linearly as following: 19 min, 8% B; 60 min, 12% B; 155 min, 35% B; 203 min, 60% B; 210 min, 75% B; 215 min, 95% B; 220 min, 95% B. Eluting peptides were directly analyzed by electrospray in an orbitrap mass spectrometer (Q-Exactive Plus, Thermo Fisher Scientific) by scanning a window of 400-2000 m/z with resolution of 70,000 at m/z 400. Tandem mass spectra were collected using HCD (32% NCE) on the 12 most intense multiple-charged parent ions at a resolution of 17,500.

Mass spectrometry data was analyzed using MaxQuant software (v.1.5.5.1) (23). Peptide identification was performed by searching against the *C. albicans* SC5314 and *C. auris* sequences from Uniprot Knowledge Base (downloaded December 6, 2017). Searching parameters included the variable modifications protein N-terminal acetylation and oxidation of methionine, in addition to carbamidomethylation of cysteine residues. Parent and fragment mass tolerance were kept as the default setting of the software. Only fully tryptic digested peptides were considered, allowing up to two missed cleaved sites per peptide. Quantification of proteins was done using the intensity-based absolute quantification (iBAQ) method (24). Intensities of each protein were normalized by the total iBAQ sum of each sample to obtain a relative number of protein copies (percentage from total). The comparison between the two species was performed by blast searches and considering a cutoff of 40% of sequence similarity to consider a protein orthologous.

### Lipid analysis

Extracted lipids were suspended in 100% methanol and analyzed by liquid chromatography tandem mass spectrometry (LC-MS/MS) as described elsewhere (25). The identification of the species was done using LIQUID software and manually inspected for validation (26). Peak intensities of each identified lipid species were extracted with MZmine v2.0 (27).

### Gas chromatography-mass spectrometry analysis

Extracted hydrophilic metabolite and lipid fractions were derivatized as described previously (28) and analyzed in an Agilent GC 7890A using an HP-5MS column (30 m × 0.25 mm × 0.25 μm; Agilent Technologies, Santa Clara, CA) coupled with a single quadrupole MSD 5975C (Agilent Technologies). The GC was set to splitless mode with the port temperature at 250 °C. Samples were injected with the oven temperature equilibrated at 60°C. The same temperature was kept for 1 minute and then raised at a 10°C/minute rate to a final temperature of 325°C for 5 minutes hold. A standard mixture of fatty acid methyl ester (FAME) (Sigma Aldrich) was used for calibrating the retention time. Retention time calibration, spectral deconvolution, and peak alignment were done with Metabolite Detector (29). Metabolites were identified by matching against FiehnLib library (30) containing additional metabolites entered in-house and/or the NIST14 GC-MS library. All identified metabolites were manually inspected.

### Quantitative analysis and data integration

Protein orthologues, lipids or metabolites were considered significantly different with a p-value ≤ 0.05 using T-test considering equal variance and two-tailed distribution. For comparative analyses, missing values were zero-filled with half of the smallest value of the dataset. Proteins were clustered by the k-means method using Multi-Experiment Viewer (MeV v4.9.0) (31), which was also used to build the heatmaps. Pathway analysis on different protein clusters was performed with DAVID (32), and specific pathways of interested were manually inspected with Vanted v2.1.1 (33). We have recently developed an R package called Rodin (https://github.com/PNNL-Comp-Mass-Spec/Rodin), to perform structural ‘lipid ontology’ (LO) enrichment analysis. A web interface Lipid-MiniOn was developed for non-R users (https://omicstools.pnnl.gov/shiny/lipid-mini-on/). Briefly, this tool creates automatically LO bins based on the lipids naming and their inferred structure, then it performs enrichment analysis using enrichment statistics to compare a Query list to an Universe (Fisher’s exact test, EASE score, Binomial test, or hypergeometric tests). In this study a Fisher’s exact test was used to perform the enrichment analysis and only the enrichments with a Fisher’s exact *p* value below 0.05 were considered to be conserved.

## Results

### Antifungal resistance

As *C. auris* is a recently-identified pathogen, its breakpoints for resistance to different antifungals have not been formally established. Given the lack of information, our results were interpreted based on the CDC breakpoint suggestions (https://www.cdc.gov/fungal/candida-auris/recommendations.html). MICs for the tested strains against amphotericin B were similar, and all strains had a MIC below 2 μg/mL, thus being susceptible against this antifungal. MMC2 was consider susceptible as the MIC to caspofungin was below 2 μg/mL. MMC1 had a MIC of 2μg/mL for caspofungin, which qualifies as resistance against this drug. Notably, *C. auris* strains were able to grow when exposed to caspofungin concentrations above their MIC, a phenomenon known as “Paradoxal effect” or “Eagle effect” (34). This effect was previously reported for *Aspergillus* and *Candida* species (34), and was very recently described for *C. auris* (35). *C. auris* MMC2 was susceptible to fluconazole, presenting a MIC at 8 μg/mL. In contrast, *C. auris* MMC1 strain was highly resistant as it was able to grow at concentrations of 1000 μg/mL of fluconazole (Table 1). As a reference, we also examined a standard *C. albicans* strain (ATCC #90028), which is susceptible to all the three drugs used in this work.

**Table 1.** Antifungal susceptibility test using the broth microdilution.

| Organism/Strain | Amphotericin B<br>(µg/mL) | Caspofungin<br>(µg/mL) | Fluconazole<br>(µg/mL) |
| --- | --- | --- | --- |
| *Candida auris* MMC1 | 1.6 | 2 | > 256* |
| *Candida auris* MMC2 | 0.8 | 1.6 | 8 |
| *Candida albicans* | 1.3 | 0.3 | 0.75 |

\*MMC1 was resistant to fluconazole concentrations of 1000 µg/mL

### Proteomic profiling of *C. auris* vs. *C. albicans*

The proteomic analysis resulted in the identification of 1869 and 2317 proteins in *C. auris* and *C. albicans,* respectively. To compare the data from these two species, we performed BLAST searches and considered orthologous proteins with more than 40% similarity. Out of the 1869 identified *C. auris* proteins, 1726 (92%) had orthologues in the *C. albicans* genome, whereas 1954 of the 2317 (84%) *C. albicans* proteins had orthologues in the *C. auris* genome. Combined, 2323 orthologues were detected in the proteomic analysis. However, only 1357 (58% of total) orthologues were consistently abundant in both *Candida* species (Table 2 and Supplemental Tables S1-S3). This indicates that despite the sequence similarity between these two species their gene expression regulation is much more divergent even in identical culturing conditions.

**Table 2.** - Identified orthologous proteins in *C. albicans* and *C. auris*

|  | *C. albicans* | *C. auris* |
| --- | --- | --- |
| Identified proteins | 2317 | 1869 |
| Non-orthologues | 363 | 143 |
| Orthologues | 1954 | 1726 |
| Total orthologues | 2323 |  |
| Orthologues present in both species | 1357 |  |

It is noteworthy that the peptides were not identical between the two species and, therefore, a quantitative proteomic analysis comparison cannot be directly achieved across the different samples. To circumvent these issues we performed an absolute quantification of each protein using the intensity-based absolute quantification (iBAQ) method and normalized each protein by a relative number of copies in the cells. The heatmap shown in Figure 1, depicts the orthologues that are differentially abundant between all three *Candida* strains. Clustering these proteins using the k-means method showed a striking similarity between the two *C. auris* isolates, but strong differences between the different species. To better understand the differences between *C. auris* strains and the *Candida* species we performed a function-enrichment analysis, which revealed that pathways such as glycolysis/gluconeogenesis, ribosomes and phagosomes were more abundant in *C. albicans*. On the other hand, *C. auris* seems to have a more active tricarboxylic acid (TCA) cycle, along with lipid and amino acid metabolism.

**Figure 1:**
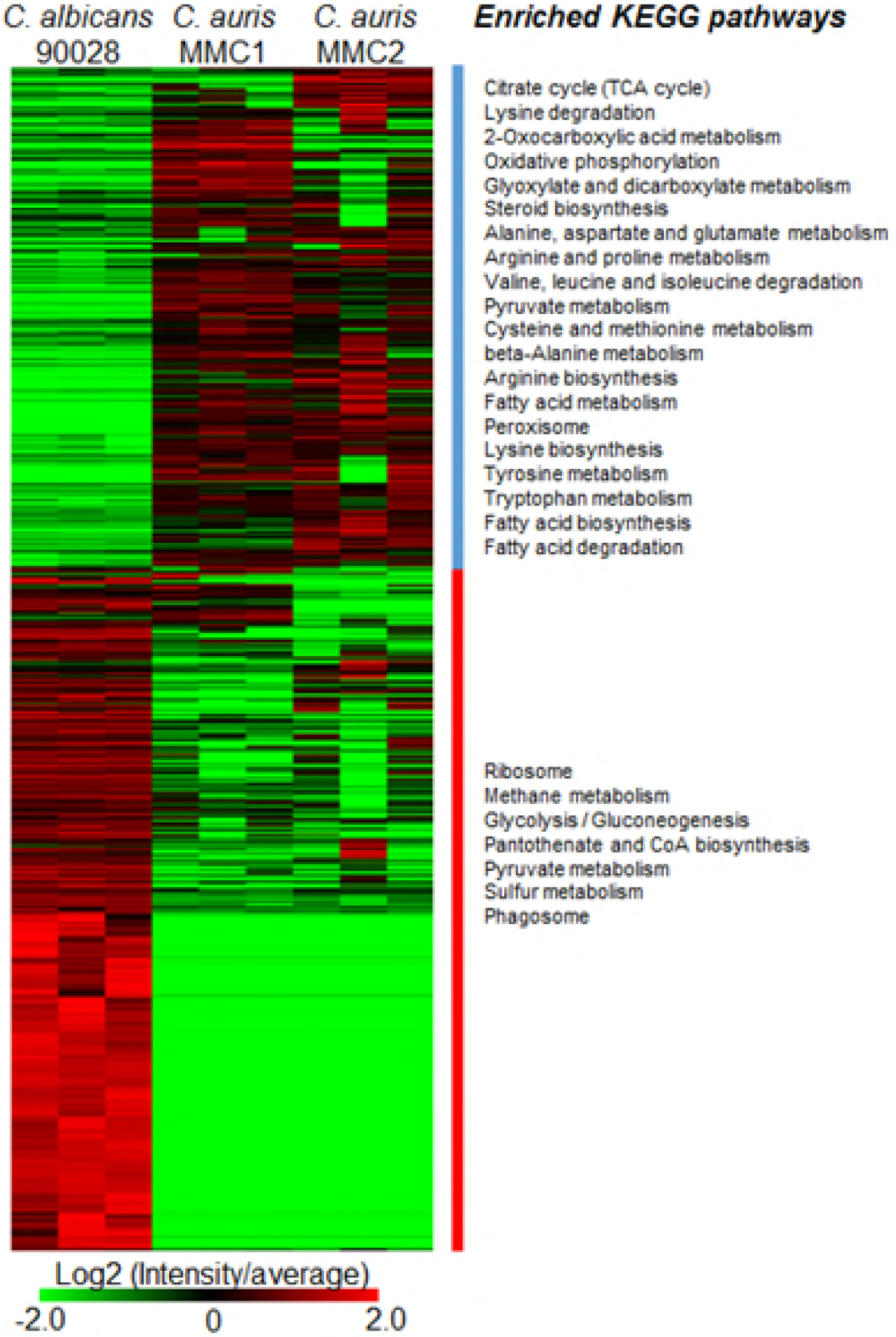
Abundance of proteins in *C. auris* and *C. albicans*. Proteins are listed in the heatmap with enriched KEGG pathways separated into two clusters based on the protein abundance between the two *Candida* species.

### Central carbon metabolism in *C. auris* and *C. albicans*

The pathway analysis showed that the glycolytic pathway was enriched in proteins with higher abundance in *C. albicans*, whereas the TCA cycle proteins were enriched with proteins more abundant in *C. auris*. Pyruvate metabolism was enriched in proteins that were more abundant in both species (Figure 1). To validate these observations and to correlate with downstream metabolic pathways, we integrated the proteomics data with a metabolite analysis into a map of central carbon metabolism. Ten out of the fifteen glycolysis/gluconeogenesis proteins were more abundant in *C. albicans* than in *C. auris*, whereas only 2 proteins were consistently more abundant in *C. auris* (Figure 2). In agreement with these observations, lactate, one of the end products of this pathway, was 16 fold more abundant in *C. albicans* than *C. auris* MMC1 and 6 fold higher than *C. auris* MMC2 (Figure 2). On the other hand, 14 out of 15 TCA cycle proteins were more abundant in *C. auris* strains than in *C. albicans* (Figure 2). Validating these observations, citrate, and fumarate had similar abundance profiles (Figure 2). In the pyruvate metabolism, proteins were not consistently more abundant in one or the other species. Some differentially abundant proteins seemed to be due to gene isoforms that were preferentially expressed between the species. For example, *C. auris* produces alcohol dehydrogenase Adh2, while *C. albicans* produces Adh5 (Figure 2). Unfortunately, the metabolites of this pathway, such as acetate, acetaldehyde, and ethanol, are small and not detectable in our GC-MS analysis. The fact that different proteins of this pathway are not uniformly more abundant in one of the species makes it more difficult to predict whether the downstream metabolic pathways would be affected. We decided to investigate the ergosterol and glycerolipids biosynthesis pathways in more detail.

**Figure 2.**
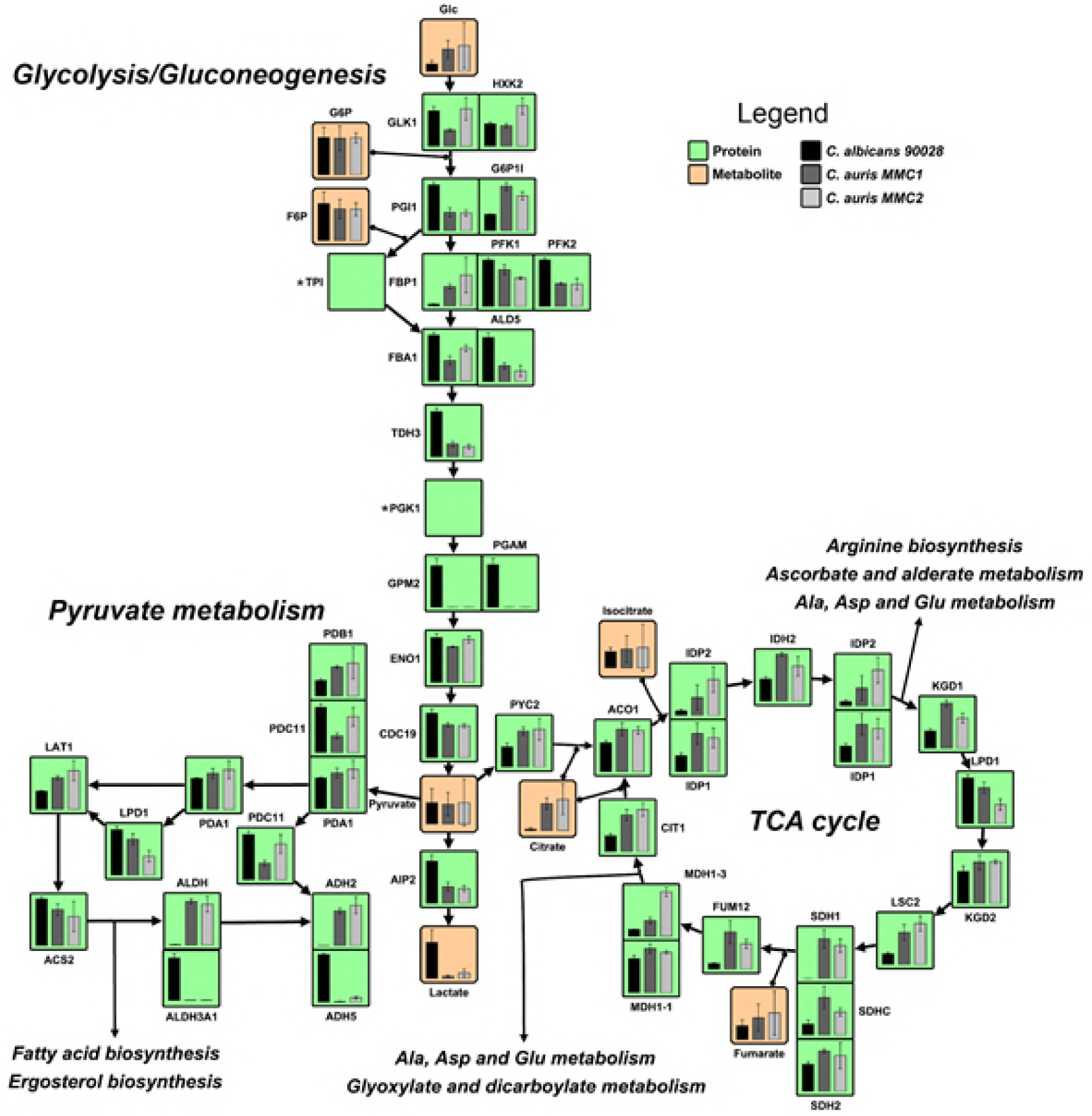
Central carbon metabolism of *C. auris* and *C. albicans*. The figure shows the relative abundance of proteins (green boxes) and the production of metabolites (orange boxes) involved in the central carbon metabolism in both *C. albicans* and *C. auris*. Paralog proteins were grouped and posted side-by-side in the map. *Genes that were only annotated in the *C. albicans* genome.

### Ergosterol biosynthesis pathway in *C. auris* vs. *C. albicans*

Fluconazole inhibits the activity of Erg11 (Lanosterol 14-alpha-demethylase), and consequently ergosterol biosynthesis. Due to the remarkable resistance displayed by MMC1 against fluconazole, we performed a comparative analysis of the enzymes and some of the metabolites present in the ergosterol synthesis pathway. Twelve (Erg10, Erg13, Erg8, Erg9, Erg1, Erg7, Erg11, Erg24, Erg27, Erg6, Erg3, and Erg5) out of nineteen of the ergosterol synthesis enzymes are more abundant in *C. auris* than in *C. albicans*, including Erg11 (Figure 3). There are, however, a few exceptions of enzymes from the ergosterol pathway that are more abundant in *C. albicans* than in *C. auris*, which is the case for Idi1 and Erg20. Farnesol, a quorum sensing molecule involved with *C. albicans* dimorphism is poorly produced by both *C. auris* strains (Figure 3).

**Figure 3.**
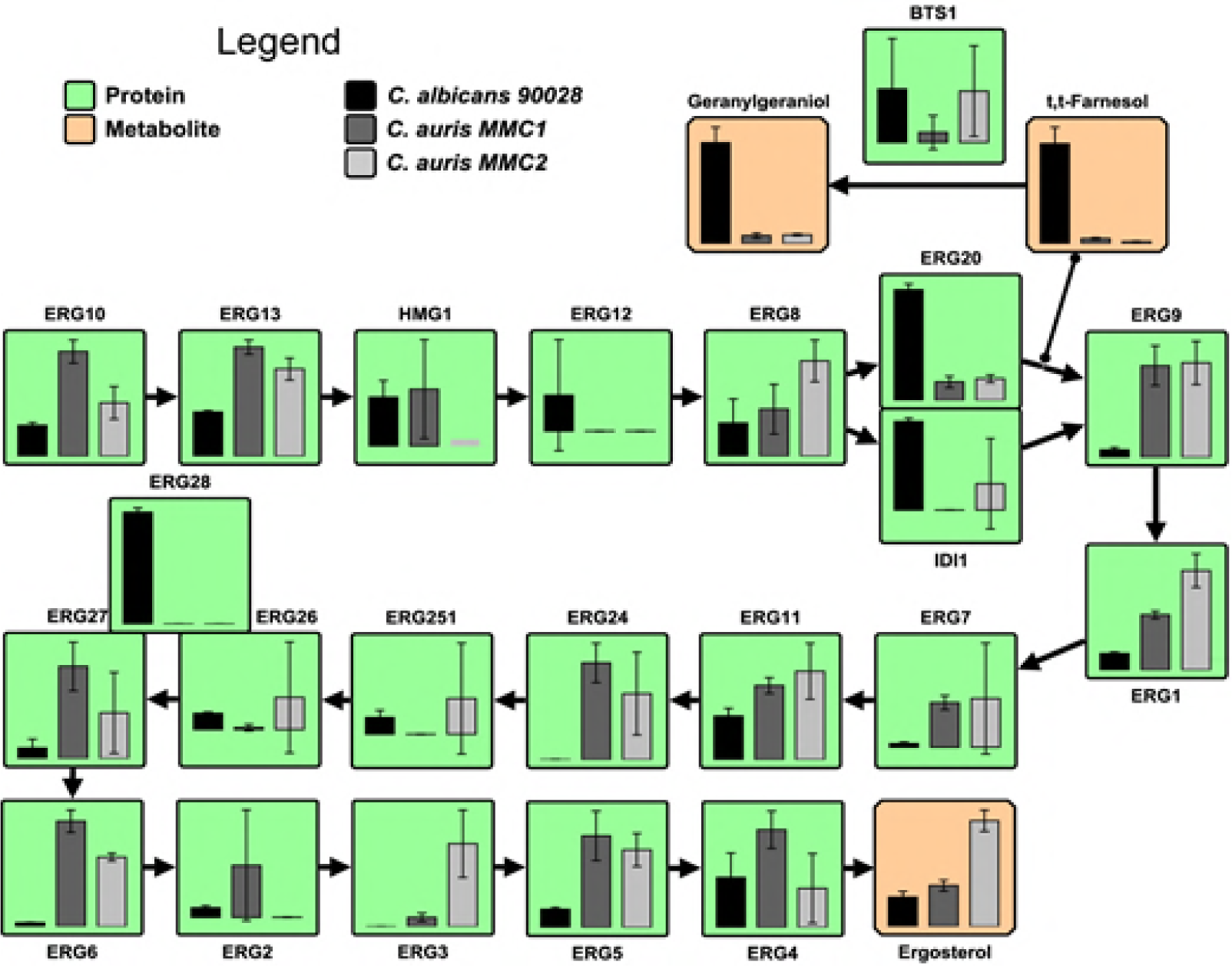
Ergosterol biosynthesis pathway in *Candida auris* and *Candida albicans*. The bar graphs represent the relative abundances of proteins (green boxes) and metabolites (orange boxes) of the pathway.

### Lipid profile of *C. auris* and *C. albicans*

The differential abundance of carbon metabolism, especially in the pyruvate metabolism, is indicative that the fatty acid biosynthesis and consequently the lipid structures could be altered. Considering that lipids are major targets of antifungal drugs (36) and part of resistance mechanisms (37, 38), we analyzed this category of biomolecules. A total of 169 lipids from 10 different classes were identified and quantified. The most diverse lipid class was triacylglycerol (TG), with 38 distinct species, followed by phosphatidylcholine (PC) with 28 (Supplemental Tables S4). To compare groups of lipids from different *Candida* species/strains, we clustered lipids based on their abundance and performed an enrichment analysis using a recently developed tool named MiniON (described in Methods). This analysis is analogous to pathway enrichment and determines whether groups of lipids are significantly enriched based on their intrinsic features (class, head group, fatty acid (FA) length and unsaturation, etc.). The results showed that TG and lipids carrying polyunsaturated FAs were enriched in *C. albicans*. Cardiolipins, lipids containing C18:3 FAs and glycerolipids carrying C16:1 FA were significantly reduced in the resistant strain MMC1 (Figure 4). Lysophospholipids were enhanced in *C. auris* MMC1 and to a less extent in *C. auris* MMC2 compared to *C. albicans*. The enriched amount of lysophospholipids is an indication of a higher phospholipase activity. We investigated the abundance profiles of enzymes with phospholipase activity in the proteomics data (Table 3). Our analysis detected seven phospholipases in *C. auris* and only five in *C. albicans*. Excepting Pld1 (A0A0L0P056), all of them were significantly more abundant in MMC1 than in *C. albicans*. Remarkably, the lysophospholipases Plb3 and Plb5 were not detected in *C. albicans*.

**Table 3.** Proteins with phospholipase activity in *C. auris* and *C. albicans*

| Protein names | *C. auris* -<br>Uniprot | *C.*<br>*albicans* | MMC1 | MMC2 |
| --- | --- | --- | --- | --- |
| Plc2p | A0A0L0P5S6 | - | ++ | + |
| Patatin-like phospholipase domain-containing protein | A0A0L0NS42 | -- | ++ | - |
| Lysophospholipase | A0A0L0NWB3 | ND | ++ | ++ |
| Lysophospholipase | A0A0L0P465 | ND | + | ++ |
| Doa1p | A0A0L0NP71 | + | ++ | ++ |
| Phospholipase | A0A0L0P056 | ++ | + | + |
| Lysophospholipase Nte1 (Intracellular phospholipase B) | A0A0L0P1C1 | ++ | - | - |

**Figure 4.**
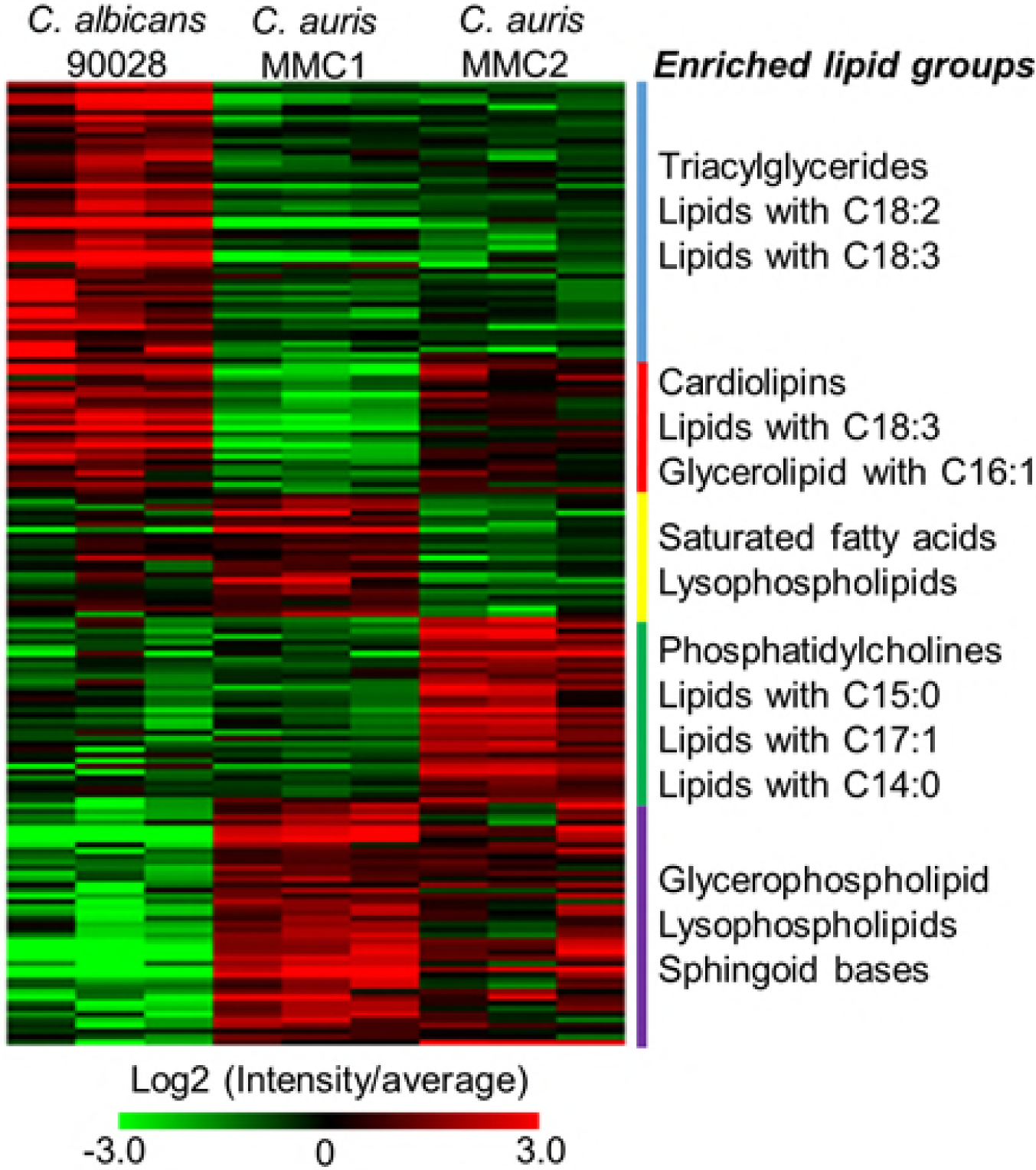
Lipid species found in *C. auris* and *C. albicans*. The abundance of all detected lipids is shown above in the heatmap. Lipids were grouped in clusters based on their abundance between different species/strains. Enrichment of lipid intrinsic features (head group, fatty acid length, fatty acid unsaturation, etc.) is listed by the side of each cluster.

*C. auris* MMC2 produced more phosphatidylcholines and lipids containing odd FAs compared to *C. auris* MMC1 and *C. albicans* (Figure 3). A GC-MS analysis of the lipid fraction indeed confirmed that C17:0 and C17:1 FAs were more abundant in *C. auris* MMC2 (Figure 5). Both strains of *C. auris* were enriched in sphingoid bases (Figure 3), which was also validated by the detection of phytosphingosine in the GC-MS analysis (Figure 4). In addition to the sphingoid bases, other sphingolipids such as ceramides, hexosylceramides and inositolphosphoceramides were also more abundant in *C. auris* MMC1 (Supplemental table S4).

**Figure 5.**
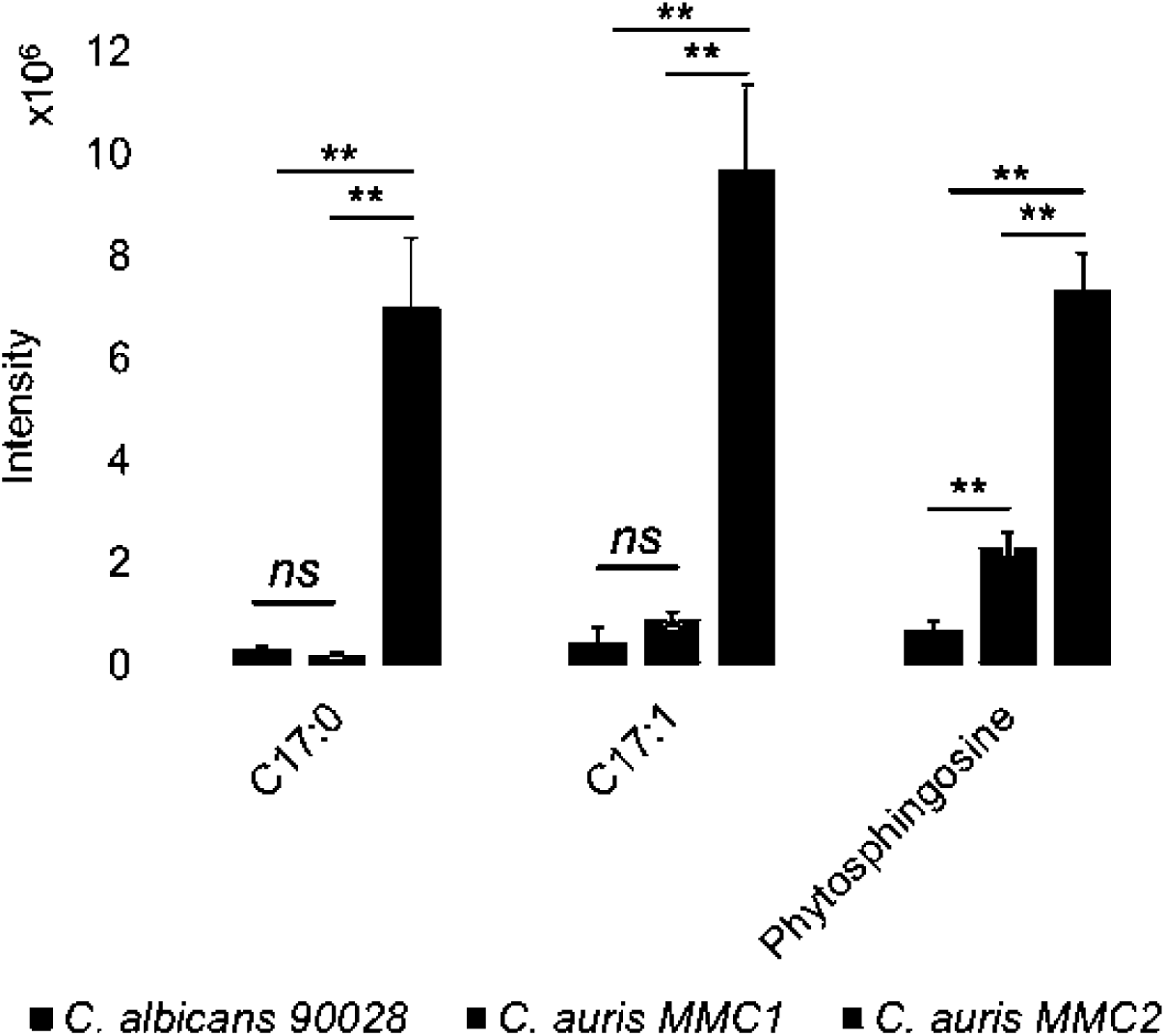
Fatty acids and sphingoid bases analyzed by GC-MS. The graph indicates the abundance of lipids containing odd FA and phytosphingosine for both *Candida* species/strains. ** p-value ≤ 0.01.

### Cell Wall Integrity (CWI) pathway, and major structural components

The proteomic analysis showed that proteins involved in the cell wall integrity (CWI) pathway displayed a significant difference between *C. albicans* and *C. auris*. Rom2, Tpk2, and the MAP kinase Mck1 were higher in strain MMC1 when compared to *C. albicans* and the fluconazole susceptible strain MMC2 (Figure 6), suggesting that the MMC1 strain is better suited to respond to antifungal drugs. Notably, the protein Pkc1 was detected only in *C. albicans*, suggesting that *C. auris* may have an alternative pathway to control CWI (Figure 6).

**Figure 6.**
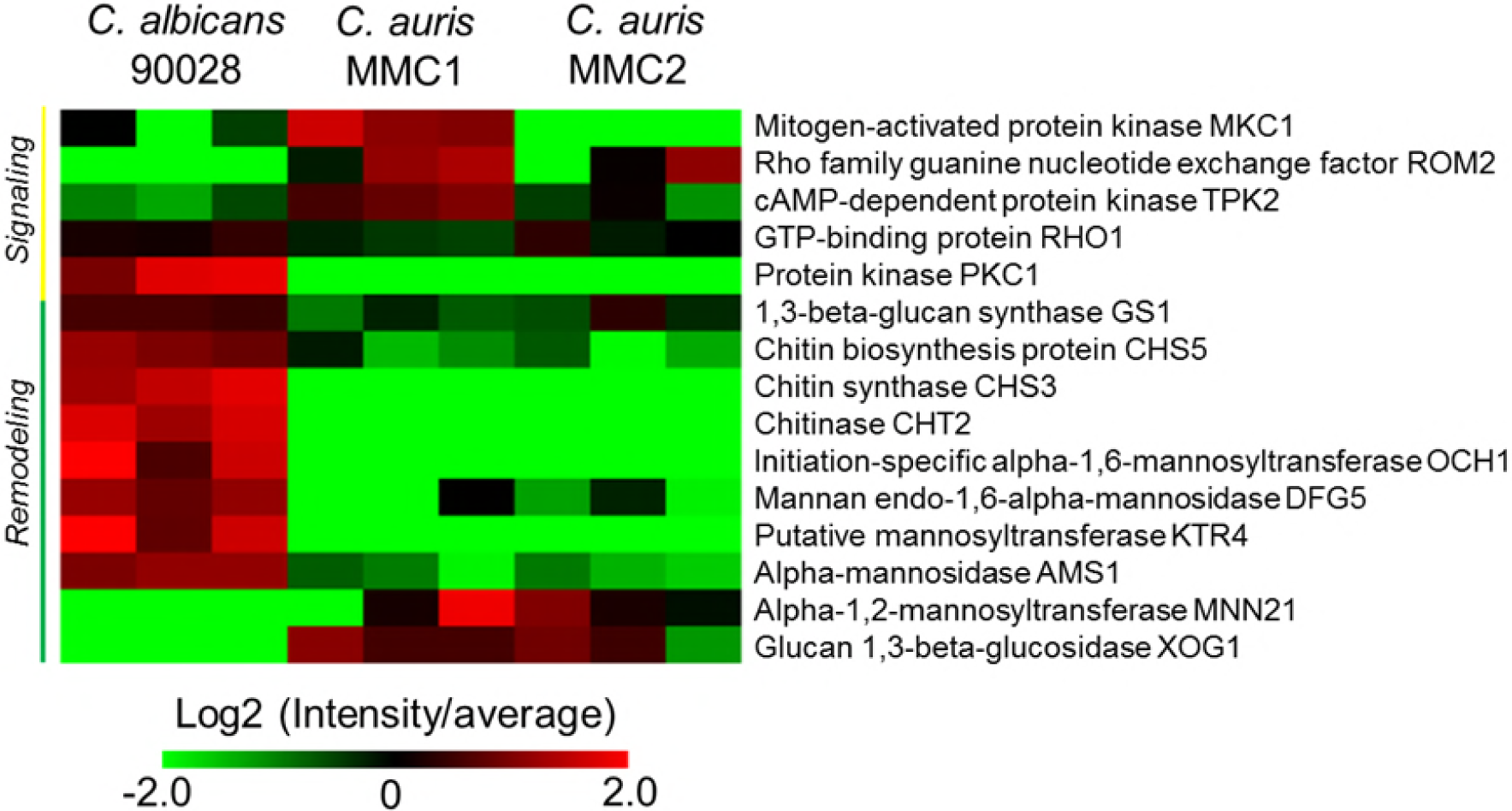
Cell wall integrity pathway. The heatmap includes signaling and major cell wall polysaccharides synthesis/degradation enzymes found in *C. auris* and *C. albicans*.

The enzymes involved in the synthesis and degradation of the major cell wall polysaccharides (glucans and chitin) and mannoproteins, were particularly distinct when *C. albicans* and *C. auris* were compared. Remarkably, chitin remodeling enzymes, β1,3 glucan synthase and most of the mannoprotein remodeling enzymes were higher in *C. albicans* when compared to both *C. auris* strains. The only exceptions were glucan 1,3-beta-glucosidase Xog1 and alpha-1,2 mannosyltransferase MN21, which were both more abundant in *C. auris* strains compared to *C. albicans* (Figure 6).

### Biofilm transcription factors and proteins

Fungal biofilms are highly resistant to drug treatment due to a combination of factors including cell density and matrix content (39). We compared the abundance of transcription factors and proteins previously reported in biofilm formation and proteins found in the biofilm matrix. Six transcription factors were reported as biofilm regulators in *C. albicans* (40-43). Our results showed that Efg1 and Ndt80 were more abundant in *C. albicans* under planktonic growth conditions with almost no abundance in *C. auris*. Remarkably, only Rob1 was more abundant in *C. auris*, specifically in the resistant strain MMC1. A list of proteins upregulated in *C. albicans* biofilms and biofilm matrix was also investigated (Supplemental Table S5). Out of 24 proteins previously reported upregulated in biofilm (44), 8 were detected in higher levels in the *C. auris* strains when compared to *C. albicans*.

### Transporters

The proteomic analysis identified 6 transporters related to drug resistance. Notably, the ABC transporter efflux pump Cdr1 and orf19.4780, an uncharacterized member of the Dha1 family of drug: proton drug antiporter, were significantly higher in the azole-resistant strain MMC1 (Figure 7). The other 4 transporters had higher abundance in other strains (Figure 7), therefore, they are less likely to be involved in the fluconazole resistance of MMC1.

**Figure 7.**
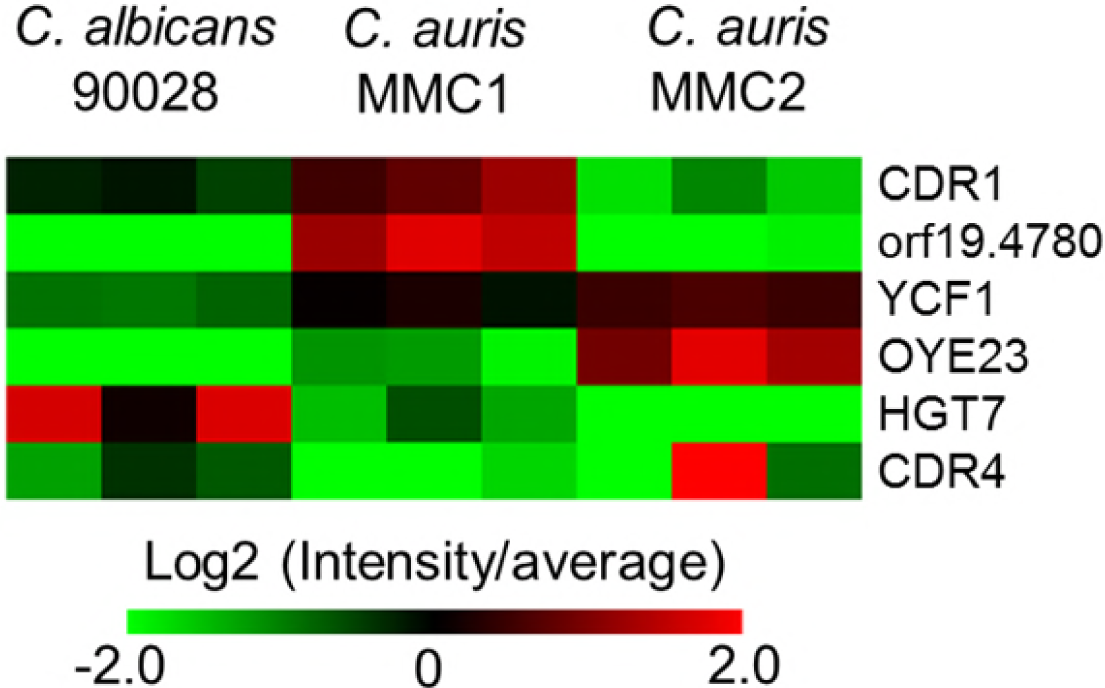
Protein abundance profile of drug resistance related transporters. The heatmap shows the detected transporters involved with drug resistance and their abundances in both *Candida* species/strains.

## Discussion

*C. auris* is an emerging pathogen that is causing extremely worrisome outbreaks across the globe. One remarkable feature of this fungus is the frequency of resistance against at least one class of antifungals. In addition, multidrug-resistant strains have been isolated from all continents. The search for a new class of antifungal drug has been a major challenge in the medical mycology community and this quest becomes even more urgent with the spread of a multidrug-resistant fungal organism like *C. auris*. In the current work, two strains of *C. auris* isolated in the Bronx, USA were analyzed by a multi-omics approach in order to better understand the molecular repertoire employed by this pathogen. In parallel to *C. auris*, we also performed the same analyses with a reference strain of *C. albicans*.

We found that MMC2 and the *C. albicans* strain were susceptible to amphotericin B, caspofungin and fluconazole, but MMC1 was resistant to both caspofungin and fluconazole. *C. auris* MMC2 MIC value of fluconazole was approximately at 8 μg/mL, which based on the CDC report, would make it a susceptible strain, even though the MIC was about 10 times higher than for *C. albicans*. Although MMC1 just met resistance criteria to caspofungin, its resistance to fluconazole was impressive, as even 1 mg/mL was not able to totally inhibit growth. The “Eagle effect”, also known as “paradoxical effect” was observed in both *C. auris* strains after treatment with caspofungin, as growth occurred at concentrations higher than the MIC.

The protein profiles from *C. auris* and *C. albicans* were qualitatively and quantitatively distinct, and both strains of *C. auris* presented very few differences from one another (Supplemental Table S3). The major observed difference between *C. auris* and *C. albicans* was in their central carbon metabolism. While proteins in the glycolysis pathway were upregulated in *C. albicans, C. auris* showed an enrichment of proteins in the TCA cycle. These results show that *C. auris* favors respiration, which is already known to be an important mechanism of fluconazole resistance in *C. albicans* by increasing ATP production and reducing oxidative stress, resulting in better overall fitness of the cell (45).

In *S. cerevisiae*, overexpression of HMG1 or deletion ERG2, can significantly increase susceptibility to fluconazole, whereas deletion of HMG1, ERG6 and ERG3, as well as overexpression of ERG11 are associated with fluconazole resistance (46). Therefore, we integrated the data of proteins and metabolites of the ergosterol biosynthesis pathway. Despite the extreme resistance of MMC1 against fluconazole, the abundance of Erg11 in this strain is similar to the observed for MMC2. On the other hand, higher abundance of Erg2 and lower abundance of Erg3 of MMC1 compared to the MMC2 isolate are in agreement with drug-resistance phenotype of MMC1. The higher abundance of Idi1 and Erg20 in *C. albicans* diverges part of the pathway to produce more isoprenoids, while *C. auris* has a more robust production of ergosterol, which is possibly involved in fluconazole resistance. Recently, sequence divergences/mutations on ERG11 in *C. auris* have been shown to be associated with resistance to azoles (47). However, the ERG11 mutations by themselves cannot explain why the level of fluconazole resistance was lower (up to 128 µg/mL) when the *C. auris* gene was expressed in *S. cerevisiae* (48). Therefore, our data combined with reports from the literature suggest that the fluconazole resistance in *C. auris* is due to modifications of multiple steps in the ergosterol biosynthesis pathway.

The lipids detected in *C. auris* were qualitatively similar to those found in *C. albicans*. However, a quantitative analysis showed that *C. albicans* has more lipids involved with energy storage, while *C. auris* has more structural glycerophospholipids and lysophospholipids. The resistant strain (MMC1) has a remarkable abundance of lysophospholipids, suggesting intense phospholipase activity. Phospholipases are virulence factors in a variety of pathogenic fungi where their activity is important for invasiveness, morphology, and persistence of infection (49-51). Phospholipase activity was recently described in *C. auris* isolates (52). In the current work, the evaluated *C. auris* strains were found to produce seven enzymes with phospholipase activity, while *C. albicans* had five of them. In addition, most of these enzymes were more abundant in *C. auris*, particularly in the resistance strain (MMC1). Corroborating these findings, an increased content of lysophospholipids was previously reported in a *C. albicans* strain adapted *in vitro* to higher concentrations of fluconazole (53). It is possible that this class of enzymes is more finely employed by *C. auris* than by *C. albicans* to promote survival and environmental adaptation for the fungus. Regarding its biological role during the host-pathogen interaction, lysophosphatidylcholine is a “find me” signal released by apoptotic cells to induce the recruitment of phagocytes to remove apoptotic bodies before an episode of secondary necrosis and enhanced inflammation (54). The MMC1 strain also had a higher abundance of sphingolipids, which can also be correlated with resistance to antifungals. These lipids are important for the assembly of membrane platforms where proteins such as drug efflux pumps are present in membrane microenvironments responsible for the export of drugs (37).

The response orchestrated by the CWI signaling pathway is central during cell wall and membrane perturbation (55). Sensors at fungal cell surface initiate a downstream cascade in order to adapt the cells under stress conditions controlling cell wall biogenesis and cell integrity (55). Remarkably, we observed that the enzymes involved with cell wall remodeling are reduced in both *C. auris* strains. However, some CWI proteins are specifically higher in the resistant strain, suggesting that the response to external signals, such as drug treatment, could be promptly controlled by the cell wall metabolism and help to explain the resistant phenotype in the MMC1 strain.

The efflux of drugs mediated by efflux pumps is an important mechanism of antifungal resistance employed by *Candida spp* (37, 56, 57). From six distinct drug efflux transporters produced by the analyzed organisms, two of them (Cdr1 and orf19.4780) were more abundant in the fluconazole-resistant *C. auris* strain (MMC1) than in the other strains. Previous publications showed that *C. auris* yeast cells, organized in a biofilm, are more resistant to antifungals than planktonic cells and correlated this phenotype with the increased expression of CDR1 (13). The impact of these efflux pumps is important during early stages of biofilm formation but decreases when it becomes mature. In mature biofilms, resistance is increased by the ability of matrix components to limit drug diffusion along with the presence of persistent cells (58). Notably, the *C. auris* strain MMC1 has a significant increase in proteins associated with biofilm formation and a higher abundance of superoxide dismutase, an enzyme involved with reactive oxygen species (ROS) detoxification and overexpressed in miconazole-tolerant persisters (58). Furthermore, a number of proteins characterized in the biofilm matrix were also higher in the resistant *C. auris* strain.

The comprehensive multi-omics approach used in this study has enabled us to begin to uncover and characterize the molecular profile of the emerging pathogen *C. auris*, which suggest a multifactorial mechanism of drug resistance in MMC1, including major differences in carbon utilization, sphingolipids, glycerolipids, sterols, cell wall and efflux pumps. Further functional omic studies that include larger numbers of *C. auris* isolates will likely have significant impact on on our understanding of the biology of this remarkable fungus and may facilitate the development of new therapeutic approaches to combat this frequently multidrug resistant yeast.

## Acknowledgments

The authors thank Erika Zink, Jeremy Teuton and Jeremy Zucker for technical assistance. Joshua D. Nosanchuk and Ernesto S. Nakayasu are partially supported by NIH R21 AI124797. Attila Gacser is supported by NKFIH K 123952, GINOP 2-3-2-15-2016-00015 and GINOP 2.3.3-15-2016-00006. Ministry of Human Capacities, Hungary grant 20391-3/2018/FEKUSTRAT is acknowledged. Parts of this work were performed in the Environmental Molecular Science Laboratory, a U.S. Department of Energy (DOE) national scientific user facility at Pacific Northwest National Laboratory (PNNL) in Richland, WA.

## Supplementary Tables

**Supplemental Table S1 -** Proteomic analysis of *C. auris* isolates. Protein abundances were normalized into relative copies numbers (see material and methods for details).

**Supplemental Table S2 -** Proteomic analysis of *C. albicans* strain 90028. Protein abundances were normalized into relative copies numbers (see material and methods for details).

**Supplemental Table S3 -** Comparative analysis of *C. albicans* strain 90028 vs. *C. auris* isolates. Protein abundances were normalized into relative copies numbers. Then values were divided by the average of the between all samples and transformed into Log2 scale (see material and methods for details). Statistically significant comparisons are highlighted in blue, while less and more abundant proteins are highlighted in green and red scales, respectively.

**Supplemental Table S4 -** Comparative lipidomic analysis of *C. albicans* strain 90028 vs. *C. auris* isolates. Lipid intensities were divided by the average of the between all samples and transformed into Log2 scale (see material and methods for details). Statistically significant comparisons are highlighted in blue, while less and more abundant lipids are highlighted in green and red scales, respectively.

**Supplemental Table S5 -** Comparative analysis of proteins from *C. albicans* strain 90028 and *C. auris* isolates involved with biofilm. Protein abundances were normalized into relative copies numbers. Then values were divided by the average of the between all samples and transformed into Log2 scale (see material and methods for details).

